# Characterization of xyloglucan-specific fucosyltransferase activity in Golgi-enriched microsomal preparations from wheat seedlings

**DOI:** 10.1101/803395

**Authors:** Richard E. Wiemels, Wei Zeng, Nan Jiang, Ahmed Faik

## Abstract

Xyloglucan (XyG) is a major hemicellulosic polymer in primary cell walls of dicotyledonous plants but represents only a minor constituent of cell walls from graminaceous monocotyledons (*Poaceae*). Our current information on XyG biosynthesis *in vitro* comes exclusively from studies on dicotyledonous plants. While XyG has been reported in grass cell walls, there are no studies of XyG biosynthesis *in vitro* in grasses. In this report, we investigated XyG structure and biosynthesis in etiolated wheat seedlings and showed that their walls contain small amounts (4-14%) of XyG. Furthermore, structural analysis using electrospray ionization mass spectrometry (ESI-MS) and high pH anion exchange chromatography (HPAEC) revealed that wheat XyG may be of XXGGG-type. Interestingly, detergent extracts from root microsomes were able to fucosylate tamarind XyG *in vitro* in a similar way as fucosyltransferase activity from *Arabidopsis thaliana* (AtFUT1) and pea (PsFUT1). Endoglucanase digestion of the [^14^C]fucosylated-tamarind XyG formed by the wheat fucosyltransferase activity released radiolabeled oligosaccharides that co-eluted with authentic fucoslyated XyG oligosaccharides (XXFG and XLFG). Although wheat fucosyltransferase activity was low, it appeared to be specific to XyG and required divalent ions (Mg^2+^ or Mn^2+^) for full activity. Together, these results suggest that the XyG fucosylation mechanism is conserved between monocots and dicots.

## Introduction

The hemicellulosic polymer, xyloglucan (XyG), makes up to 35% of the dry weight of primary cell walls of dicotyledonous plants [1–4]. Cell walls from graminaceous monocotyledons (*Poaceae*) contain only 4-10% XyG, and this XyG is structurally less complex than XyG from dicots [2;5;6]. All XyGs consist of a backbone of β-(1,4)-linked glucosyl (Glc) residues, and depending on the substitution pattern of this backbone with α (1,6)-linked xylosyl (Xyl) residues, XyG structure was initially classified into two types, namely XXXG- and XXGG-type [7], where X and G designate substituted and unsubstituted Glc residues in the backbone, respectively (for nomenclature see [8;9]). Other XyG types, such as the XXX-type, have been identified in the coats of *Helipterum eximium* seeds [10], the XXGGG-type in species of the subclass Asteridae [11], and the XXXXG-type in cotyledons of *Hymenaea courbaril* [12]. Based on analytical data from XyG purified from immature barley plants [13] and rice seedlings [14], XXGGG-type was proposed for XyG from grasses with glucan backbone regions having more or less substitution [6;14]. However, the presence and purification of the XXGGG repeating unit itself was not demonstrated.

Side-chain substitutions on XyG have been shown to be structurally diverse and species/tissues/development-dependent. Although it was suggested that Xyl residues of XyG from grasses may occasionally bear terminal galactose (Gal) or arabinofuranose (Ara*f*) residues [6;15;16], this was never experimentally confirmed through purification of the polymer. XyG from Solanaceae is of XXGG-type, and it has been shown that it can be further arabinosylated and/or galactosylated [17]. Several works demonstrated that cell walls of monocots contain fucosylated XyGs [18–20]. In some species, the backbone glucosyl residue can be *O*-acetylated [11;17;21]. The non-xylosylated glucosyl backbone residues of the XXGG-type oligosaccharides are often *O*-acetylated [17], which is not reported in XXXG-type XyGs [22]. According to recent structural studies at least 24 unique, naturally occurring xyloglucan side-chain structures exist [9;22;23]. However, linking this structural diversity of XyG substitutions to physiological functions is still elusive. The Arabidopsis *xlt2 mur3.1* double mutant, in which XyG is lacking galactosyl, fucosyl, and acetyl residues showed a dwarfed plant [24;25].

Until now, the biochemistry of XyG biosynthesis *in vitro* has been investigated exclusively in dicots. For example, biochemical studies using microsomal membranes from dicotyledonous plants showed that the incorporation of Glc and Xyl residues into the “xylosyl-glucose” backbone occurs simultaneously, which requires a synergistic/cooperative mechanism between XyG-glucan synthase (XGSase) and XyG-xylosyltransferase (XXT) activities [26–29]. This mechanism has not been confirmed yet in any membrane preparation from monocotyledonous plants. Many Arabidopsis (*Arabidopsis thaliana*) glycosyltransferase (GT) genes involved in XyG biosynthesis have been identified and characterized, providing target genes with which to investigate XyG biosynthesis in other plant species including economically important plants such as wheat, rice, and maize. These genes include a *XyG-fucosyltransferase* (*AtFUT1/MUR2*) from the CAZy GT37 family [30], several *XyG-xylosyltransferases* (*XXTs*) from the GT34 family [31–35], and two *XyG-galactosyltransferases* (*MUR3* and *XLT2*) from the GT47 family [24;36]. MUR3 and XLT2 are the galactosyltransferases for addition of Gal to the first and second Xyl residues, respectively, from the reducing of an XXXG unit. The β-(1,4) glucan backbone is synthesized by a XGSase that belongs to cellulose synthase-like C (CSL-C) family [37].

One of the major differences between XyGs from dicots and grasses is fucose (Fuc) content. Although there is evidence that fucosylate XyG is present in monocots [6;14;18;19;38;39], and a fucosyltransferase gene has been identified in rice (*Oryza sativa*) genome (using functional complementation approach [20]), biochemical characterization of fucosyltransferase activity from any plant tissues of the *Poaceae* is lacking. Here, we present evidence of fucosyltransferase activity in microsomal preparation from etiolated wheat seedlings. Interestingly, detergent-solubilized activity from microsomal membranes of root tissues is able to fucosylate tamarind XyG *in vitro* in a similar way as Arabidopsis AtFUT1 and Pea PsFUT1 enzymes. Structural analysis of XyG oligosaccharides (XyGOs), released by treatment with a XyG-specific endoglucanase, suggested that cell walls of wheat roots may contain XyG of two types: XXGGG- and XXXG-type. Although we show that these XyG types contain Fuc, Gal, and Ara residues, we were not able to determine whether they constitute different domains of the same polymer or are part of two distinct polymers synthesized by different machineries. To our knowledge this work represents the first report on the biochemical characterization of a XyG fucosyltransferase activity in grasses, and thereby furthers our understanding of XyG biosynthesis in monocots.

## Results

### Xyloglucan from etiolated wheat seedling walls contains Fuc, Gal, and Ara residues

We reasoned that if XyG from wheat seedlings are fucosylated, we should be able to detect the presence of oligosaccharides containing Fuc and Gal in KOH extracts of wheat cell walls. Thus, alcohol insoluble residues (AIR) prepared from cell walls of 6-day old etiolated wheat seedlings were directly treated with 4M KOH to extract the non-cellulosic polymers, and XyGOs were released by enzymatic treatment of these extracts with a purified XyG-specific endoglucanase (XG5, [40]). Typically, 4M KOH solubilized ∼25% and ∼33% (w/w, based on phenol-sulfuric assay) of the wall material from roots and shoots, respectively. Treatment of these alkali extracts with XG5 released ∼13% and ∼46% (w/w, based on phenol-sulfuric assay) of the material, respectively. Control reactions containing boiled XG5 enzyme released less than 0.5% of the material. These results allowed us to roughly estimate XyG contents in wheat root and shoot to ∼4 and ∼14% (w/w), respectively.

Next, the endoglucanase-released material was analyzed for monosaccharide composition through total acid hydrolysis followed by HPAEC fractionation. Purified XyG from pea treated in the same conditions and same enzyme was used as a positive control. As expected, Glc and Xyl were the two main monosaccharides in all XyGOs (pea and wheat) and Gal, Ara, and Fuc were also present, but at lower amounts (Table 1). According to our estimates, Gal and Fuc contents represent 5 mol% and <0.5 mol% in root oligosaccharide mixtures, respectively (Table 1), suggesting that only a small fraction of the Gal is fucosylated. The identity of Fuc from wheat XyGOs was further confirmed using gas chromatography-mass spectrometry (GC-MS) analysis (data not shown). The Glc:Xyl ratio in wheat XyG from roots and shoots is 2.19 and 2.27, respectively; compared to a mixture of pea XyG oligosaccharides (released with XG5 treatment) from which has a Glc:Xyl ratio of 1.22 (Table 1). As a control, analysis of pea XyGOs generated the expected ratios Glc:Xyl:Gal:Fuc of 7.5:6:1.7:1, respectively, which are typical of primary wall XyGs from dicots [1;41]. However, despite purification, XG5 endoglucanase was still contaminated with β-glucan hydrolase activity (lichenase, see below), which can act on β(1,3)(1,4)-mixed-linkage glucan (MLG, an abundant polymer in cell walls of grasses) that explain the overestimated Glc amount in wheat XyGOs. In conclusion, monosaccharide analysis suggests the presence of Fuc in XyGOs from wheat root but not detectable in XyGOs from shoots. Also, the detection of Ara in this analysis would suggest the presence of an arabinosylated side chain in wheat XyG.

**Table 1:** Monosaccharide composition of wheat XyG oligosaccharides (XyGOs) released from KOH-extracts of wheat cell walls by digestion with XyG-specific endoglucanase (XG5 from *Aspergillus aculeatus*). KOH-extracts were prepared from root and coleoptile cell walls. Released wheat XyG Oligosaccharides were hydrolyzed with 2M TFA (120°C, 1h) and the released monosaccharides were desalted and then analyzed by HPAEC on a CarboPac PA20 column (Dionex). The values are the averages of triplicate analyses plus/minus standard deviation (SE). ND stands for not detected. Pea XyG oligosaccharides were prepared from purified XyG that was treated with XG5.

|  | Wheat roots | Wheat shoots | Purified pea XyG |
| --- | --- | --- | --- |
|  | *(mol %)* |  |  |
| Glucose | 60 ±2 | 63 ±2 | 45±2 |
| Xylose | 26 ±1 | 28 ±1 | 37 ±1 |
| Arabinose | 8 ±1 | 6 ±1 | Traces |
| Galactose | 5 ±1 | 3 ±1 | 10 ±1 |
| Fucose | 0.5 ±0.1 | ND | 6 ±1 |
| Glucose:xylose ratio | 2.30 | 2.25 | 1.22 |
| Glc:Xyl:Gal:Fuc ratio | 120:52:16:1 | 21:9:1:0 | 7.5:6:1.7:1 |

We next analyzed wheat XyGOs by electrospray ionization mass spectrometry (ESI-MS) and ESI-MS/MS. As indicated in Figure 1, the XG5-released XyGOs showed signal ions that are characteristic of XyGs, namely ions at mass/charge ratio *m/z* 792, *m/z* 953, *m/z* 1085, *m/z* 1115, and *m/z* 1247 (all as sodiated ions [M+Na]^+^) corresponding to oligosaccharides having 3 hexoses (Hex) and 2 pentoses (Hex3Pen2), Hex4Pen2, Hex4Pen3, Hex5Pen2, and Hex5Pen3, respectively. Importantly, all the ions detected were absent in control reactions where boiled XG5 enzyme was used (Fig. 1C). Purified XyGOs from etiolated pea seedlings were included as a positive control and, as expected, the most abundant oligosaccharides were XXXG, XLXG/XXLG, and XXFG corresponding to ions at *m/z* 1085, 1247, and 1393 (all as [M+Na]^+^), respectively (Fig. 1D). The ions at *m/z* 792 (Hex3Pen2), *m/z* 953 (Hex4Pen2), and *m/z* 1115 (Hex5Pen2), which were observed in wheat XyGOs but not in pea XyGOs, suggest that at least a part of the glucan backbone of wheat XyGs is less substituted with Xyl residues. On the other hand, the ions at *m/z* 1085 (Hex4Pen3) and *m/z* 1247 (Hex5Pen3), which were present in both wheat and pea XyGOs, may indicate that some regions of the backbone may contain “Gal-Xyl” (L) and/or “Ara-Xyl” (S) side chains (Fig. 1). We also consistently observed ions at *m/z* 1013 and 1175 that correspond to oligosaccharides having six hexoses (Hex6) and seven hexoses (Hex7), respectively, supporting the presence of MLG oligosaccharides due to lichenase activity contamination in our XG5 preparation, despite our efforts to purify XG5 activity from the original enzyme preparation. This was further confirmed by testing our purified XG5 preparation on MLG from barely, which released gluco-oligosaccharides including Hex6 and Hex7 oligosaccharides (data not shown).

**Figure 1:**
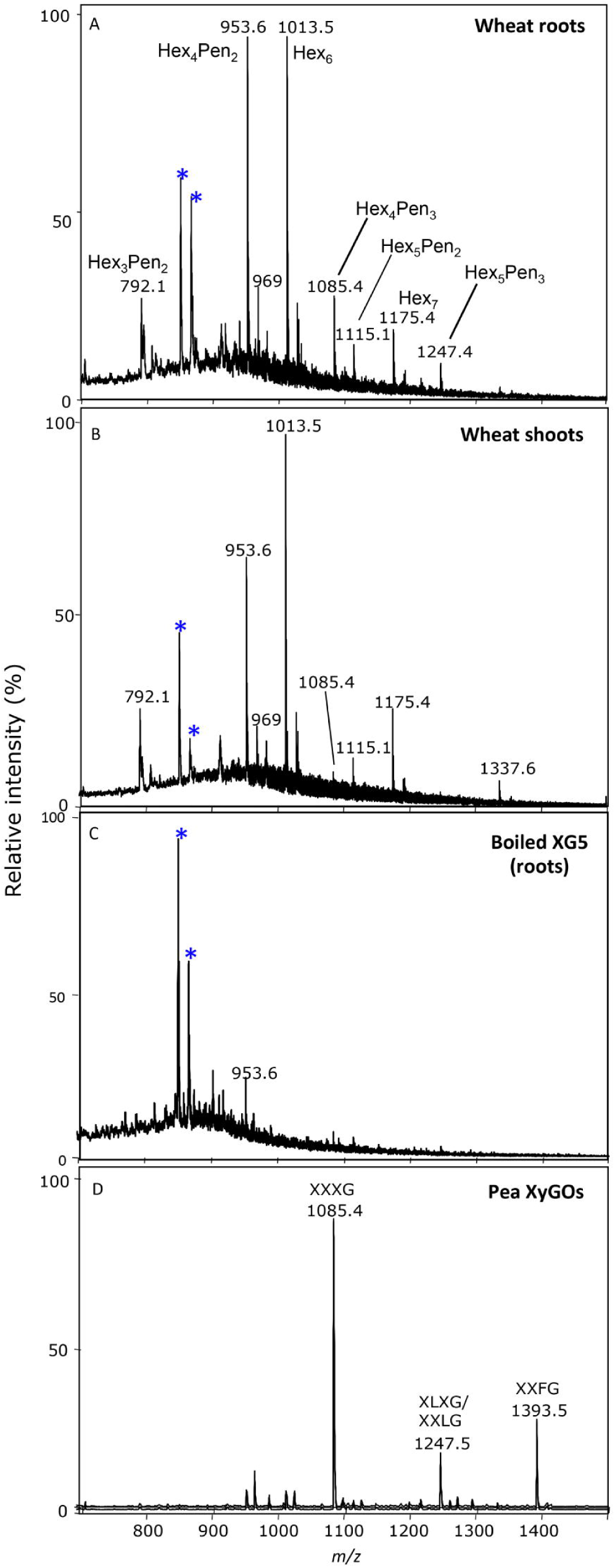
Electrospray ionization mass spectrometry (ESI-MS) analysis of XyG oligosaccharides (XyGOs) from root and shoots (coleoptile) walls of 6-day-old etiolated wheat seedlings. XyGOs were released from KOH-extracted polymers by treatment with purified XyG-specific endoglucanase XG5 from *Aspergillus aculeatus*. Panel A, Typical ESI-MS spectrum of XyGOs from roots. Panel B, Typical ESI-MS spectrum of XyGOs from shoots. Panel C, Typical ESI-MS spectrum of solubilized material from incubation of boiled XG5 enzyme with KOH extracts from wheat roots. Panel D, Typical ESI-MS of purified pea XyGOs used as controls. Pea XyG was digested with purified XG5 and XyGOs released were purified on Bio-gel-P2 column. * Indicates contaminant impurities from the enzyme or KOH extracts.

To gain more insights about the structures of the wheat XyGOs, their fragmentation patterns was analyzed by collision-induced dissociation (CID)-MS/MS). CID-MS/MS fragmentation induces the cleavage of both glycosidic bonds and bonds within the sugar ring [42;43]. The cleavage of glycosidic bonds usually results in the loss of either the whole sugar molecule (e.g. 180Da for Glc) from the reducing end, or the sugar molecule minus water (e.g. 162Da for Glc) from the non-reducing end [42]. However, cleavage of a sugar ring yields so-called cross-ring fragments as a result of a loss of the acidic residue (^k,l^A_n_ ions, see [42] for nomenclature). Furthermore, XG5 endoglucanase only cleaves XyG backbone between an unbranched and a branched Glc residues [40]. Based on this mode of action, the end products of XG5 should be XyGOs with one or more unbranched Glc residues at their reducing ends and no unbranched Glc residues at their non-reducing ends. CID-MS/MS fragmentation analysis of the ion at *m/z* 792 (Hex3Pen2, [M+Na]^+^) depicted in Figure 2A showed the presence of four abundant product ions formed by loss of 18, 60, 120, and 132Da from the parent ion. Loss of 60 and 120Da were the result of cross-ring fragmentation of the Glc residue at the reducing end, indicating that most Hex3Pen2 oligosaccharides have at least one unbranched Glc at their reducing ends. Loss of 132Da (Xyl minus water molecule) resulted in the production of the ion at *m/z* 659 (Fig. 2A). This fragmentation pattern supports the presence of XXG fragments among the Hex3Pen2 oligosaccharides. Similarly, CID-MS/MS analysis of the ion at *m/z* 953 (Hex4Pen2, [M+Na]^+^) resulted in cross-ring fragmentation of the Glc residue at the reducing end, as indicated by the three abundant ion signals corresponding to the loss of 18, 60, and 120Da from the parent ion (Fig. 2B). The ion signals at *m/z* 644 and 660, which are the result of the loss of a disaccharide “Hex-Pen” from the non-reducing end, and the ion at *m/z* 791 are due to the loss of a disaccharide “Hex-Hex” from the reducing end. Based on the mode of action of XG5 and the fragmentation pattern of the parent ion, XXGG is the most abundant oligosaccharide in the Hex4Pen2 XyGO mixture. Furthermore, the ion signals at *m/z* 701, 731, and 761 resulting from cross-ring fragmentation of the second Glc residue at the reducing end provide additional support to the presence of two Glc residues at the reducing end (Fig. 2B).

**Figure 2:**
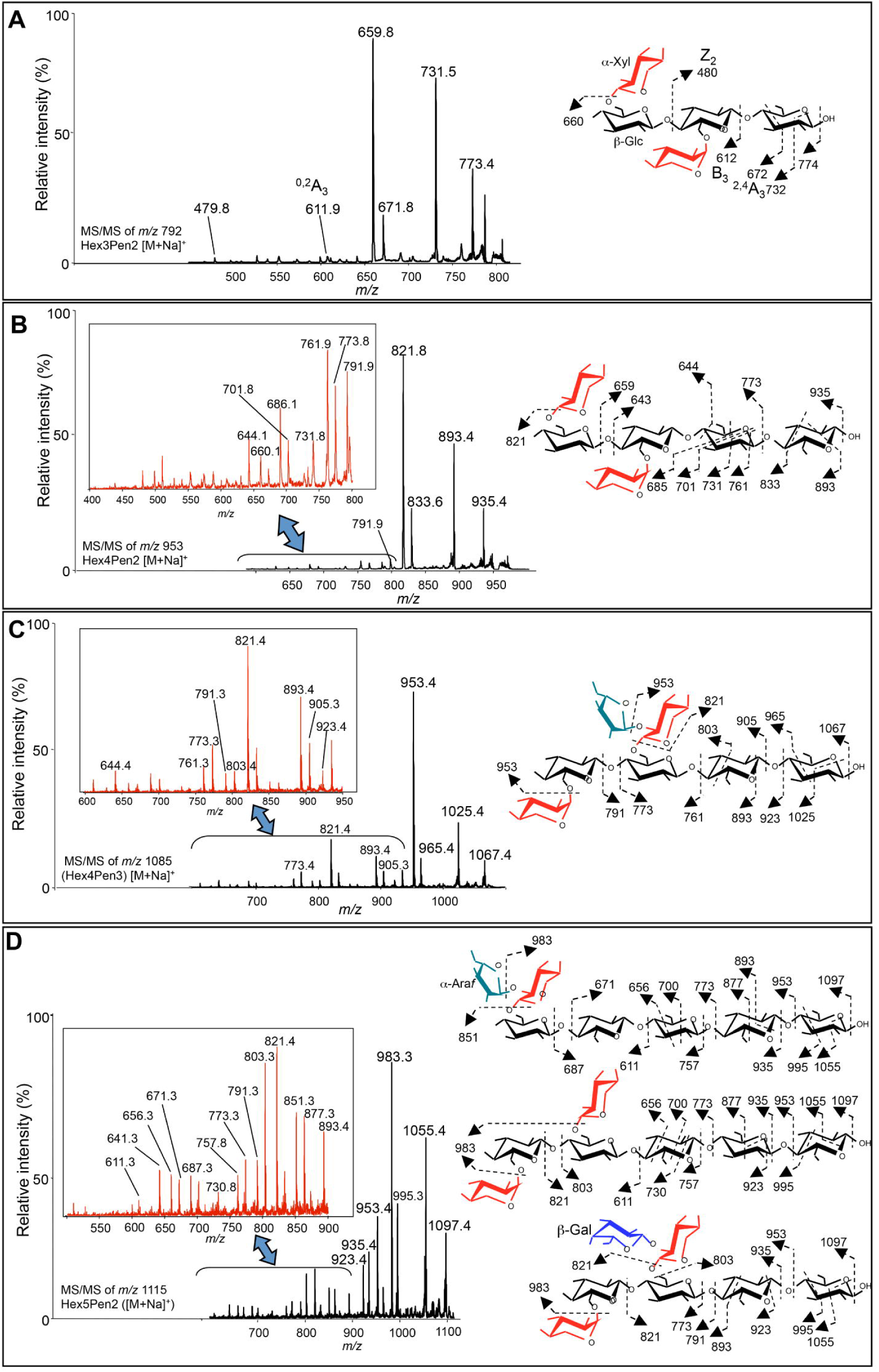
Collision-induced dissociation mass spectrometry analysis (CID-MS/MS) of XyGOs from wheat roots (see Fig. 1, Panel A). Typical CID-MS/MS spectra of ions at *m/z* 792 Hex3Pen2 (Panel A), *m/z* 953 Hex4Pen2 (Panel B), *m/z* 1085 Hex4Pen3 (Panel C), and *m/z* 1115 Hex5Pen2 (Panel D). Structures of some oligosaccharides deduced from MS/MS data are shown with fragmentation scheme on the right of each panel. Note the presence of “arabinosyl-xylose” and “galactosyl-xylose” side chains.

Fragmentation of the ion at *m/z* 1085 (Hex4Pen3, [M+Na]^+^) also resulted in cross-ring fragmentation ions at the Glc residue of the reducing end (*m/z* 1067, 1025, and 965, Fig. 1C). While the ion signals at *m/z* 773 and 791 are due to a loss of a “Xyl-Glc” disaccharide from the non-reducing end, the ions at *m/z* 905 and 923 correspond to the loss of two Hex residues (i.e., “Glc-Glc” disaccharide) from the reducing end. Importantly, the presence of the ion at *m/z* 821, which can be formed only if two pentoses are lost at the same time (Fig. 1C), is an indication of XyGOs may have “Xyl-Xyl” (U) and/or “Ara-Xyl” (S) side chains. Therefore, the most likely structure of this oligosaccharide is XSGG (Fig. 2C).

Fragmentation of the ion *m/z* 1115 ([M+Na]^+^) representing Hex5Pen2 oligosaccharides showed a more complex spectrum, suggesting a mixture of XyGO structures such as XXGGG, XLGG/LXGG, and SGGGG. Most of the ion signals are due to cross-ring fragmentation of two or three unbranched Glc at the reducing end (Fig. 2D). The loss of the trisaccharide “Gal-Xyl-Glc” from the non-reducing end resulted in the formation of a weaker ion signal at *m/z* 641 and supports the presence of an LXGG structure (Fig. 2D). Similarly, the ion signal at *m/z* 851, corresponding to an oligosaccharide containing five Hex residues is a result of the loss of “Ara-Xyl” disaccharide (S side-chain) from the non-reducing end of SGGGG (Fig. 2D). The presence of this oligosaccharide is also supported by the ion signals at *m/z* 671 and 687, which are due to the loss of “Ara-Xyl-Glc” trisaccharide from its non-reducing end (Fig. 2D).

CID-MS/MS spectrum of the ion *m/z* 1247 (Hex5Pen3, [M+Na]^+^) also showed a complex spectrum (data not shown) that was difficult to interpret. However, careful analysis of this spectrum suggested a mixture of XyGOs having structures similar to *m/z* 1115 with an additional pentose (most likely Xyl) forming the following possible XyGO structures: SXGGG/XSGGG, XXXGG, XLXG/XXLG, and SLGG. This means either that XyG from wheat roots has XXXG and XLXG/XXLG structures (found in dicot plants) or these tissues produce a polymer that has both 4-unit backbone repeats and 5-unit backbone repeats. Because of the low content of Fuc in wheat XyG, we were not able to identify fucosylated XyGOs or purify oligosaccharides containing Fuc and/or Gal. Further work is required to determine the fine structure of wheat XyG(s). Only when successful purification of wheat XyG polymer is achieved can the fine structure be determined.

### Microsomal fractions from roots but not shoots of etiolated wheat seedlings contain XyG-fucosyltransferase activity with similar biochemical characteristics as AtFUT1 and PsFUT1 activity

The presence of fucosylated XyG in cell walls of etiolated wheat seedlings suggests that a XyG-specific fucosyltransferase activity is present in these tissues. To investigate this hypothesis, Triton X100-solubilized proteins from Golgi-enriched microsomes from etiolated wheat seedlings (coleoptile and root) were tested for their ability to transfer [^14^C]Fuc from GDP-[^14^C]Fuc onto tamarind XyG in a standard XyG fucosyltransferase assay [44]. As indicated in Table 2, substantial fucosyltransferase activity was observed in wheat root tissues, which is in agreement with the fact that Fuc was detected in XyGOs from cell walls of wheat seedling root. However, the wheat XyG-fucosyltransferase specific activity (∼30 pmol Fuc incorporated per hour per mg protein) was ∼10-times less than was detected in equivalent detergent solubilized extracts from pea microsomes used as a positive control (Table 2). This activity was only detected in roots, as Triton X-100 extracts from wheat coleoptiles (shoots) did not transfer any radioactivity onto tamarind XyG (Table 2). Time course analysis of wheat XyG-fucosyltransferase activity from detergent extracts of microsomes prepared from roots harvested at various developmental stages, namely five, eight, and 11 days after germination, indicated that tissues from 5-day old wheat seedlings have the highest XyG-fucosyltransferase activity (Fig. 3A), suggesting that the activity is developmentally regulated. This is consistent with results from studies of XyG-fucosyltransferase activities in pea and Arabidopsis where higher activities were observed in young and rapidly dividing tissues [30;36]. XyG-fucosyltransferase activity was also detected in the roots of etiolated Brachypodium seedlings, though the activity was 10 times lower than in wheat (data not shown). Furthermore, wheat fucosyltransferase activity is specific for XyG, as other cell wall polymers such as rhamnogalacturonan-I (RG-I), known to contain terminal α-fucosyl residues, [45], arabinan, and pectic galactan were not good acceptor substrate in the assay (Fig. 3C).

**Figure 3:**
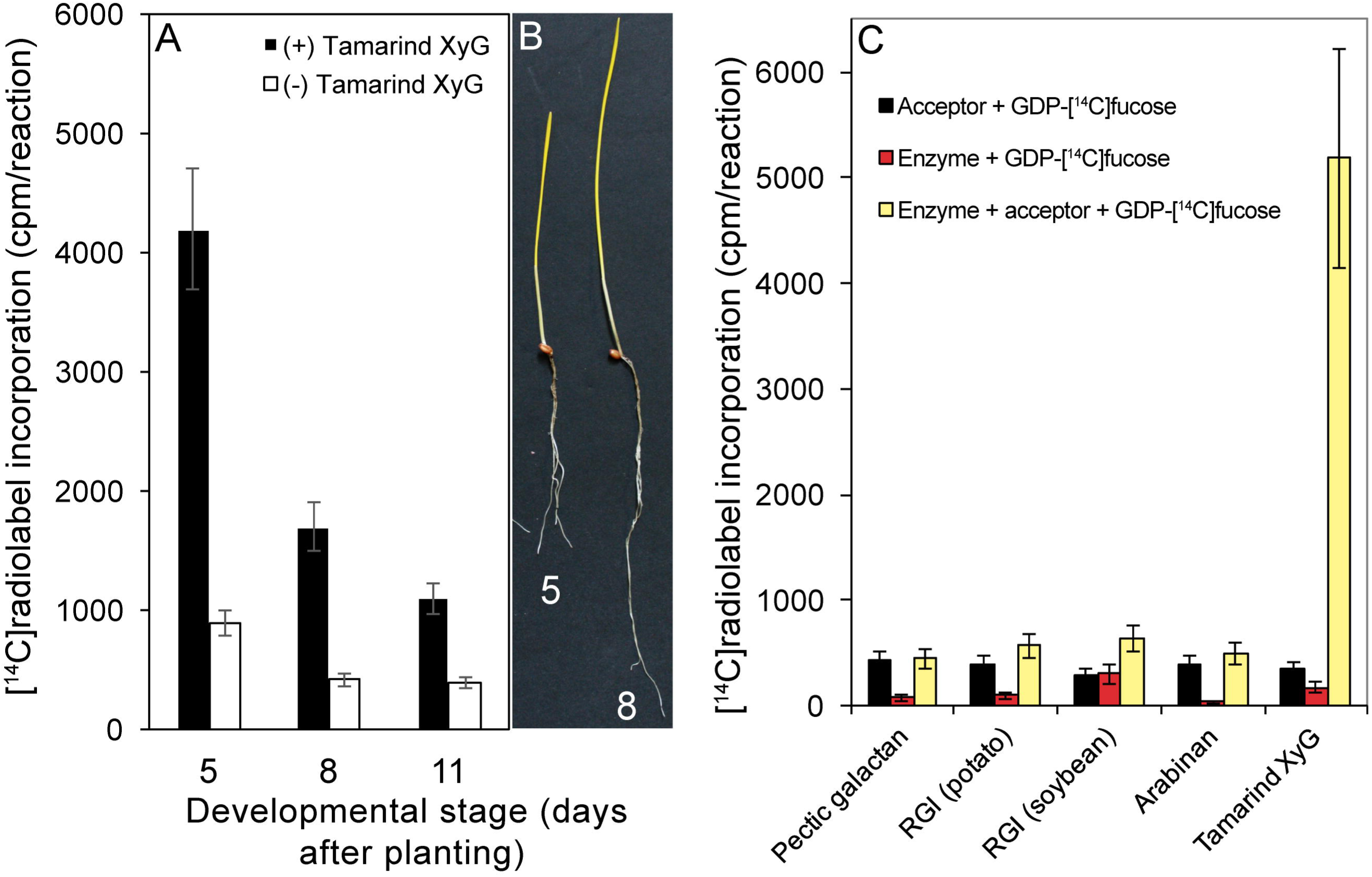
Xyloglucan-fucosyltransferase activity in Golgi-enriched microsomal membranes from wheat roots. Panel A, Wheat activity in Triton-X100-solubilized proteins from microsomal membranes obtained from roots of five-, eight-, and 11-day-old etiolated wheat seedlings. The activity is measured as the amount of [^14^C]Fuc transfer onto tamarind XyG (expressed as cpm/reaction). Reactions lacking tamarind XyG (acceptor) are used as negative controls. Panel B, Picture of five- and eight-day-old etiolated wheat seedlings given as a reference. Panel C, Specificity of the wheat activity tested on several cell wall polymers as substrate acceptors (*i.e*, arabinan, pectic galactan, RGI from potato, RGI from soybean) compared to tamarin XyG using Triton-X100-solubilized proteins. Reactions lacking either the acceptor or the enzyme are included to show that [^14^C]Fuc incorporation is XyG-dependent. Error bars in panels A and C represent the SE of at least two biological replicates.

**Table 2:** Transfer of [^14^C]radiolabeled fucose ([^14^C]Fuc) from GDP-[^14^C]Fuc to tamarind XyG by Triton X-100-extracts from microsomes of etiolated wheat and pea seedlings. Detergent-soluble proteins from wheat (0.2mg/reaction) or pea (0.36mg/reaction) were incubated with 100µg tamarind XyG and GDP-[^14^C]fucose (65,000cpm) for 1h at room temperature, and the reactions were stopped by precipitation with 1mL 70% (v/v) ethanol. [^14^C]radiolabel incorporation (expressed as pmol fucose/h/mg protein) was measured as described in Materials and Methods section. The values are from experiments repeated at least five times. Deviations represent standard deviation (SE).

| Tissues | [$^{14}\text{C}$]Fuc incorporation | |
| --- | --- | --- |
|  | No tamarind XyG | Plus tamarind XyG |
|  | *(pmol/h/mg protein)* |  |
| Wheat coleoptile | 10.3±1.5 | 11.3±2.5 |
| Wheat root | 3.4±1.5 | 31±1 |
| Pea hypocotyl | 8.5 ±2 | 306 ±10 |

In a second experiment, we investigated the metal ion dependence of wheat XyG-fucosyltransferase activity in the presence and absence of various divalent cations. Wheat XyG-fucosyltransferase activity does not require divalent cations for activity; however, the addition of 5mM MgCl_2_ more than doubled activity, and addition of CaCl_2_ enhanced activity by ∼40% and MnCl_2_ had no effect (Fig. 4A), which is consistent with the previous study on XyG-fucosyltransferase activity in pea [29]. Wheat XyG-fucosyltransferase activity is optimal at ∼pH 6 (Fig. 4B), which is also consistent with pea XyG-fucosyltransferase activity [29]. Finally, wheat activity is specific for GDP-Fuc because GDP-mannose and GDP-Glc did not produce any product with tamarind XyG (data not shown).

**Figure 4.**
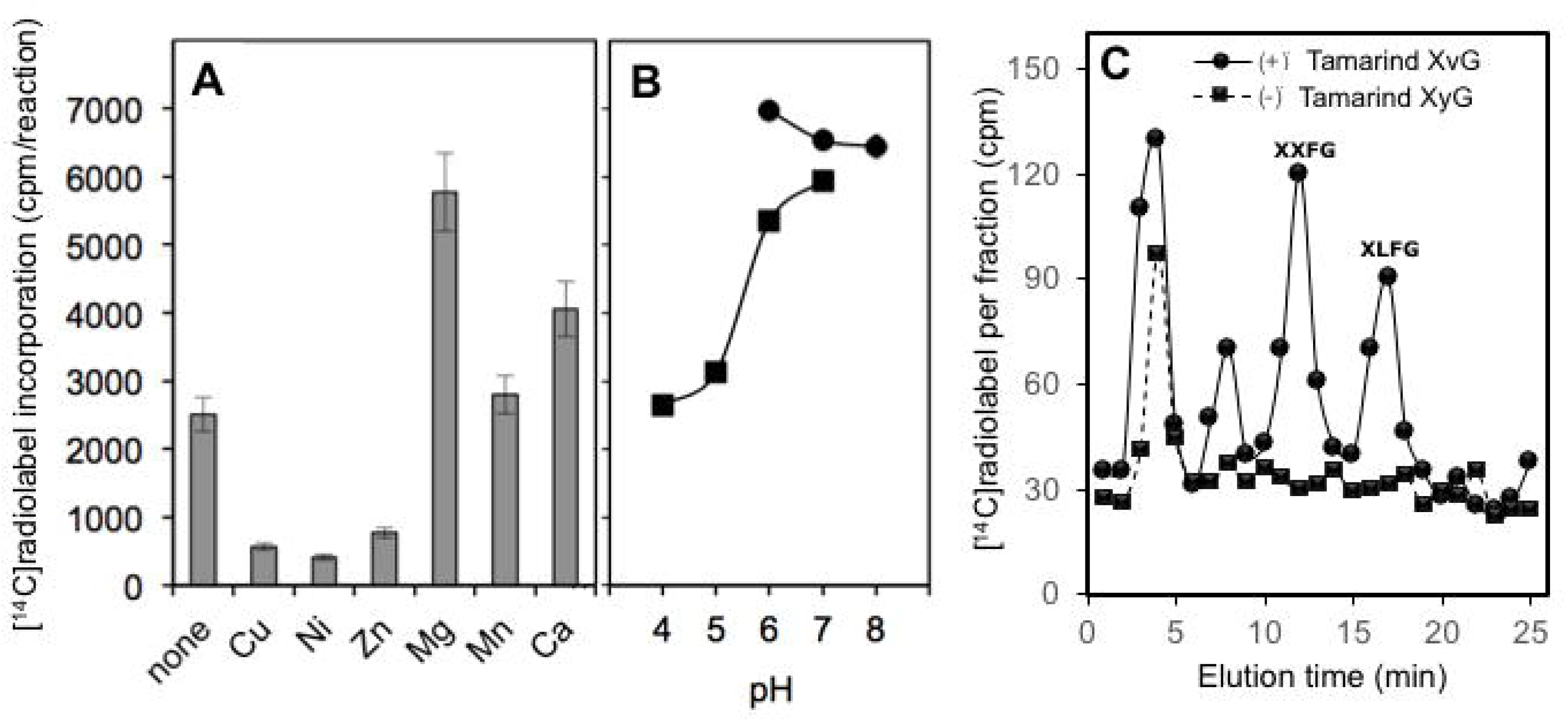
Effect of divalent ions (Panel A) and pH (Panel B) on wheat XyG-fucosyltransferase enzyme activity and analysis of its radiolabeled products by HPAEC (Panel C). XyG-fucosyltransferase activity was measured in Triton-X100-solubilized proteins from pea microsomal membranes (reaction conditions are provided in Materials and Methods). [^14^C]fucose-tamarind XyG product was generated by incubating detergent-solubilized proteins from wheat root microsomes in the presence of GDP-[^14^C]fucose and tamarind XyG, and then digested with endoglucanase (Megazyme). The released [^14^C]fucosylated tamarind XyGOs were fractionated by HPAEC on a CarboPac PA200 column (Panel C). Control reactions containing no tamarind XyG (▪) are compared to the assay containing tamarind XyG as substrate acceptor (•). Elution time of known fucosylated XyGOs (XXFG, XLFG) used as standards are indicated at the top of the peaks. Error bars in panel A represent the SE of at least two biological replicates.

### Wheat enzyme activity fucosylates tamarind XyG in a similar way as pea enzyme activity

Next, we wanted to verify whether wheat XyG-fucosyltransferase activity fucosylates tamarind XyG in a similar manner as the activity from pea [29]. Thus, the product “[^14^C]Fuc-tamarind XyG” generated by the wheat microsomal extracts was treated with an endoglucanase (E-CELTR, Megazyme) and the released [^14^C]radiolabeled fragments were fractionated by high pH anion exchange chromatography (HPAEC). Figure 4C shows a typical elution profile of these fragments and indicates that over 50% of the [^14^C]radiolabel co-eluted with two authentic fucosylated XyGOs, XXFG and XLFG, generated and purified from pea XyG. Around 25% of the [^14^C]radiolabel eluted at ∼4min and may be attributed to free [^14^C]Fuc. Around 18% of the [^14^C]radiolabel eluted as smaller XyGOs with unknown structure at around eight minutes (Fig. 4C). However, previous work showed that XFG, LFG, and FG usually elute in this time range on CarboPac-PA-100 column [46]. The [^14^C]products from a control reaction (lacking tamarind XyG) were digested and analyzed under the same conditions. As indicated in Figure 4C, the three main peaks (XXFG, XLFG, and unknown XyGOs) were absent, and the peak at 2-3min was present. These data support the conclusion that the detergent-solubilized wheat activity from microsomal membranes of etiolated wheat have a XyG-dependent fucosyltransferase activity that can fucosylate tamarind XyG to generate a polymer that contains XXFG and XLFG oligosaccharides. In addition, we found that the incorporation of [^14^C]Fuc was greater with tamarind XyG as an acceptor compared to XyG from nasturtium seed (data not shown), which might be explained by the difference in the fine structure of these two storage XyGs. Tamarind XyG has relatively more XXLG subunits compared to nasturtium XyG [47]; XXLG was shown to be a better acceptor for the fucosyltransfer reaction in pea [29]. There was no sign of products fucosylated closer to the non-reducing end of the subunits, e.g. XFXG or XFLG. We would expect these subunits to elute differently from one another but closer to XXFG and XLFG. No peak corresponding to labeled XFFG was observed, which would elute from this column at a different locus from XXFG and XLFG, as it does after HPLC on a Dynamax-60A NH_2_ column [48]. Thus, we conclude that wheat XyG-fucosyltransferase activity is specific for fucosylation of galactosyl residues nearest the reducing end of XyG subunits.

### Identification of wheat members of the GT37 family

To gain further insights into fucosylation of XyG in wheat seedlings, we sought to identify putative wheat fucosyltransferases (FUTs) in public genomics resources (Table 3). As of June 2019, CAZy database lists only one wheat member (CAMPLR22A2D_LOCUS5750). Using bioinformatics approach (see Materials and Methods for detailed approach), we identified 16 wheat members of the GT37 family (not including homeologs), a number that is comparable to rice whose genome contains 17 GT37 members. These wheat FUTs were named TaFUT-A through TaFUT-P (Table 4). Of the 16 wheat members, only four were full-length (*TaFUT-A*, *TaFUT-B*, *TaFUT-C*, and *TaFUT-H*), and three wheat genes (*TaFUT-D*, *TaFUT-F*, and *TaFUT-O*, Table 4) were cloned in this study. To facilitate the identification of wheat genes that cluster with the two known XyG-fucosyltransferases at the time of performing this work: AtFUT1 and PsFUT1 (from pea, *Pisum sativum* L.), we carried out phylogenetic analysis using Arabidopsis, rice and wheat members of the GT37 family. According to phylogenetic analysis, wheat orthologs of all rice genes were identified, except for Os08g0334900 (Fig. 5A). Rice and wheat FUTs clustered into two major groups (II and III) and a smaller group (group IV). Group II and III can be further split into two subgroups (A and B) that cluster in the same branch as FUTs from dicots (Fig. 5A). Phylogenetic analysis also showed that FUTs from dicots (Arabidopsis and Pea) clustered together in a clade (group I) that contains no rice or wheat FUTs (Fig. 5A). Thus, the phylogenetic analysis could not resolve the relationship between the known XyG-fucosyltransferases (AtFUT1 and PsFUT1) and putative rice and wheat FUTs. Even pairing of AtFUT1 and PsFUT1 (both are from two dicot species) was not possible (Fig. 5A). It seems that the FUTs have independently diverged in each plant lineage, which makes identification of grass orthologs of characterized FUTs not possible using phylogeny. Although, group IV (TaFUT-O and Os02g0764400) are phylogenetically the most closely related to AtUT1 and PsFUT1 (Fig. 5A), pairwise alignments of the full-length putative grass FUTs in groups II-A and II-B (8 putative wheat FUTs) with Arabidopsis FUTs (group I) showed high sequence identity/similarity with AtFUT1 and PsFUT1 compared to any other Arabidopsis FUT sequence. This makes the selection of wheat candidates for testing even more challenging. TaFUT-O showed 54% identity at the amino acid level (67-69% similarity) with PsFUT1 and AtFUT1. Thus, *TaFUT-O* was chosen for enzyme assay testing. We also selected arbitrary TaFUT-F and TaFUT-D from group II-B.

**Figure 5:**
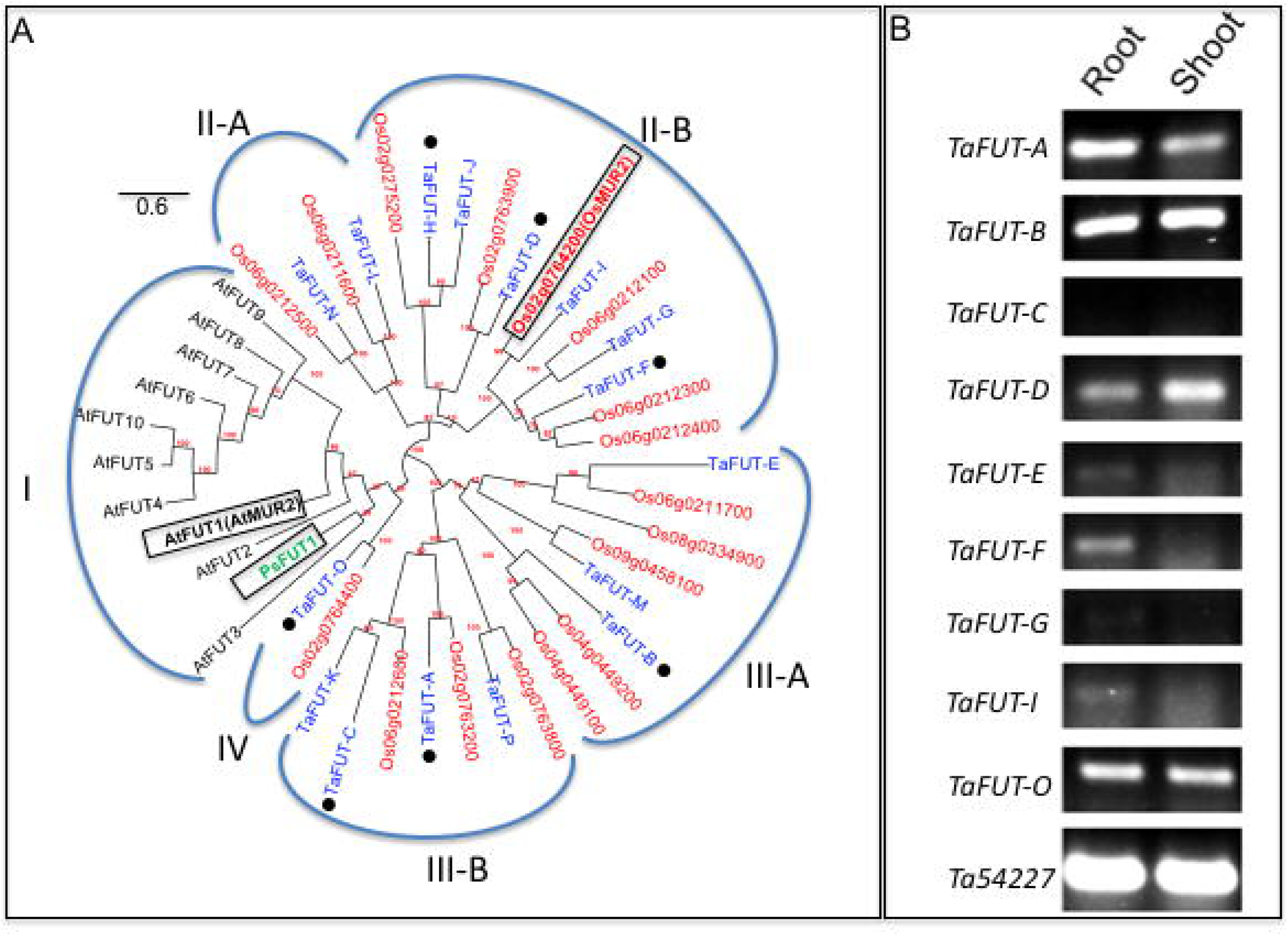
Phylogenetic and expression analyses of wheat members of the GT37 family. Panel A, Phylogenetic analysis of Arabidopsis (black, At-), wheat (blue, Ta-), and rice (red, Os-) members of GT37 family. Pea FUT1 (green) is also included. The tree was constructed with protein sequences using Phylogeny.fr platform as described in “Materials and Methods.” Branch support (to infer bootstrap values) is based on an approximation of the standard Likelihood Ratio Test. Panel B, RT-PCR for nine *TaFUT* genes using cDNA prepared from total RNA from root and shoot from five-day-old etiolated wheat seedlings. The control gene, *Ta54227*, encodes for a AAA-family member of ATPases [79].

**Table 3:**
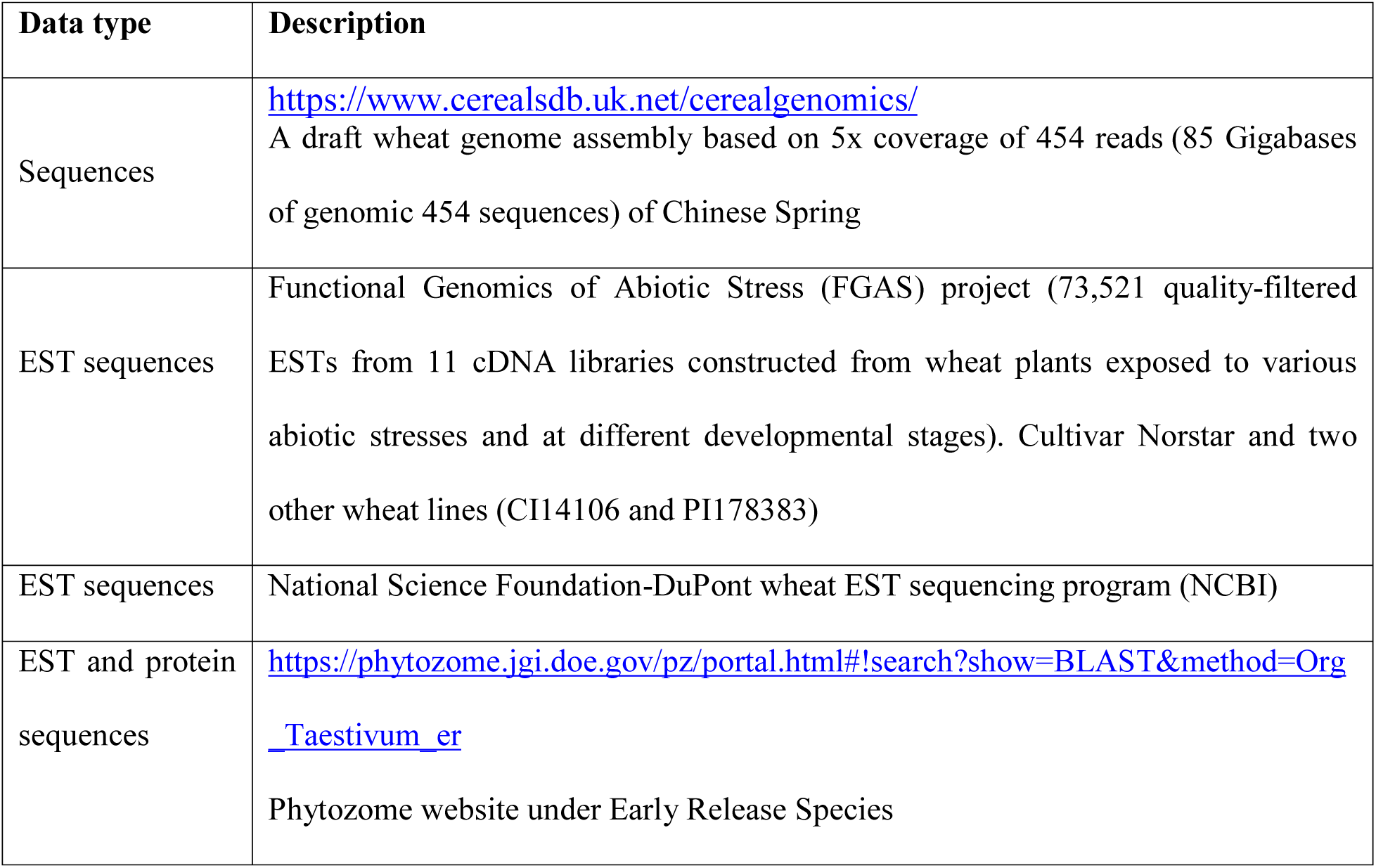
Data available in the wheat Database.

| Data type | Description |
| --- | --- |
| Sequences | https://www.cerealsdb.uk.net/cerealgenomics/<br>A draft wheat genome assembly based on 5x coverage of 454 reads (85 Gigabases of genomic 454 sequences) of Chinese Spring |
| EST sequences | Functional Genomics of Abiotic Stress (FGAS) project (73,521 quality-filtered ESTs from 11 cDNA libraries constructed from wheat plants exposed to various abiotic stresses and at different developmental stages). Cultivar Norstar and two other wheat lines (CI14106 and PI178383) |
| EST sequences | National Science Foundation-DuPont wheat EST sequencing program (NCBI) |
| EST and protein sequences | https://phytozome.jgi.doe.gov/pz/portal.html#!search?show=BLAST&method=Org_Taestivum_er<br>Phytozome website under Early Release Species |

**Table 4:** List of wheat members of CAZy GT37 family identified by our bioinformatics approach (see Materials and Methods for details) and found in Phytozome database. Rice homologs are also listed. The gene highlighted in red were cloned in this work.

| Identified by our approach (including homeologs) | ID in Phytozome (including homeologs) | Closest rice homolog | Predicted size (amino acids) | Full-length |
| --- | --- | --- | --- | --- |
| TaFUT-A-1<br>TaFUT-A-2<br>TaFUT-A-3 | Not present | Os02g0763200 | 576 | Yes |
| TaFUT-B | Traes_2AL_07BFC11CA.1 | Os04g0449100 | 544 | Yes |
| TaFUT-C | Traes_4DS_37A53FCC2.1<br>Traes_4BS_233289BB6.1 | Os06g0212600 | 561 | Yes |
| **TaFUT-D** | Traes_6DL_E724AD52E.2<br>Traes_6BL_2981FD934.1 | Os02g0763900 | 570 | Yes |
| TaFUT-E | Traes_7DS_854E2AD57.2 | Os06g0211700 | 521 | No |
| **TaFUT-F** | Not present | Os06g0212300<br>Os06g0212400 | 589 | Yes |
| TaFUT-G | Not present | Os06g0212100 | 279 | No |
| TaFUT-H | Traes_2AL_5A666899B.1 | Os02g0275200 | 563 | Yes |
| TaFUT-I | Traes_6DL_B04B4C48D.1 | Os02g0764200<br>(OsMUR2) | 214 | No |
| TaFUT-J | Traes_1BL_B0FD45910.1 | Os02g0275200 | 449 | No |
| TaFUT-K | Not present | Os06g0212600 | 501 | No |
| TaFUT-L | Not present | Os06g0211600 | 513 | No |
| TaFUT-M | Traes_5BL_9812A4B62.9 | Os09g0458100 | 494 | No |
| TaFUT-N | Not present | Os06g0212500 | 489 | No |
| **TaFUT-O** | Traes_6AL_D0DAF923E.1<br>Traes_6BL_F1AC805AA.2<br>Traes_6DL_E31E7DEE8.1 | Os02g0764400 | 555 | Yes |
| TaFUT-P | Traes_6DL_1334C2E40.1 | Os02g0763800 | 344 | No |
| Not identified | Not present | Os08g0334900 |  |  |

Since XyG-fucosyltransferase activity is detected in roots, we reasoned that its encoding gene must be highly expressed in roots compared to shoots. Thus, in a second step, we sought to determine the levels of expression of wheat FUT genes in roots and shoots using RT-PCR technique. For this experiment, we focused on 8 wheat FUT genes (*TaFUT-A*, *TaFUT-B*, *TaFUT-C*, *TaFUT-D*, *TaFUT-E*, *TaFUT-F, TaFUT-G*, *TaFUT-I*, and *TaFUT-O*), since they were either full-length or have enough long nucleic sequence to design gene-specific primers. Our RT-PCR results summarized in Figure 5B shows that *TaFUT-A*, *TaFUT-B*, *TaFUT-D*, and *TaFUT-O* are expressed in roots and shoots at almost equal levels, while *TaFUT-F* and *TaFUT-I* are expressed predominantly in roots (with TaFUT-I having very low expression level). *TaFUT-C* transcript was not detectable in both roots and shoots (Fig. 5B). Thus, according to expression data, either or both TaFUT-F and TaFUT-I could be XyG-fucosyltransferases. By the time this work was completed, a rice XyG-fucosyltransferase gene (*Os02g0764400*) was identified through complementation of Arabidopsis *axy2.2* mutant [20]. While Os02g0764400, TaFUT-F, and TaFUT-I clustered together in group II-B, TaFUT-I is the closest homolog to Os02g0764400 (both located in the same branch, Fig. 5A). At the time of performing this work, TaFUT-I (Traes_6DL_B04B4C48D.1, Table 3) was not among the 13 wheat FUT sequences publicly available. It was identified after the recent early release of species in Phytozome v12.1.6. In addition, expression data indicate that *TaFUT-I* has very low expression levels in roots. Therefore, we did not include TaFUT-I in our original work and focused on TaFUT-F and TaFUT-O for enzyme activity testing.

His-tagged versions of TaFUT-F and TaFUT-O were produced in Pichia cells and detergent-solubilized proteins were prepared from independent transgenic yeast cell lines, each expressing one of these putative FUT genes. Expression of these putative FUTs was confirmed by western blot analysis using the 6xHis antibody. Detergent-solubilized proteins from microsomal membranes were used in the FUT assay as described in [44]. Unfortunately, none of the detergent extracts showed XyG-dependent activity (data not shown). More work is needed to identify wheat XyG-fucosyltransferase gene.

## Discussion

Etiolated seedlings have been studied extensively in many plants, including cereals, as they represent relatively simple and homogenous tissues with active primary cell wall metabolism. The growth of the shoots (coleoptiles) in these etiolated seedlings occurs mainly through rapid and intensive elongation of cells and cell wall material synthesis [6;49;50]. Thus, etiolated seedlings are a good model to investigate polysaccharide biosynthesis in primary cell walls. Using this plant system, we showed that detergent-solubilized extracts from Golgi-enriched microsomal membranes from roots were able to fucosylate tamarind XyG *in vitro*. This activity was higher in younger seedlings when elongation of cells and cell wall synthesis are at their maxima. This finding has an important implication as it indicates that wheat XyG contains some “L” side chains because a XyG-fucosyltransferase activity *in vivo* in wheat can only be expected if the galactosylated accepter contains “L” side chains (*i.e*., XXLG, XLGG, or possibly XLGGG). However, such a galactosylated substrate has never been unambiguously and directly demonstrated in XyG from *Poaceae*. Based on linkage data, Sims et al. [16] suggested that side chains in XyGs from *Poaceae* might contain Gal and Ara residues. However, the authors conceded that further studies would be necessary to confirm such structures. Our data is in agreement with previous work showing evidence that monocots cell walls contain fucosylate XyG [6;14;18;19;39;40;51]. Using ESI-MS analysis of XyGOs released from AIR preparation by treatment with purified XyG-specific endoglucanase, XG5 [40], we showed that XyG contents in wheat root and coleoptile (shoots) walls were roughly estimated at ∼4 and ∼14%, respectively. These levels are comparable to published data in other cereals such as barley seedlings [6;52;53] suspension-cultured maize cells [54], rice endosperm [55], and rice seedlings [14]. The observed difference in XyG content between coleoptile and root walls may be explained by a difference in XyG biosynthesis and turnover, which are usually associated with rapid cell expansion in dicots, and a similar observation was also reported for some graminaceous monocots [56]. In the case of etiolated wheat seedlings, shoot cells undertake extensive and rapid expansion (compared to root) during growth, which may explain their 3.5-times more XyG content compared to root walls. Although the walls of wheat shoots contain more XyG compared to root walls, XyG-fucosylatransferase activity was mostly confined to roots. Recent analysis of XyGs in rice showed that fucosylated XyG is also confined to young root tissues [20]. The presence of fucosylated XyG in other grasses, such as miscanthus, foxtail millet, and rice, was indirectly demonstrated using specific antibodies for fucosylated XyG (i.e., CCRC-M1 and CCRC-M106) [57]. Further analysis of wheat shoot XyGOs released by XG5 treatment using CID-MS/MS revealed the presence of oligosaccharides that can only be attributed to a XyG of the XXGGG-type. These XyGOs seem to come from XyG domains of limited or no substitution (e.g. XXGGG and SGGGG) and domains with greater substitution (e.g. XXXG and XLGG) that contain “L” side chain. The presence of an ion at *m/z* 1247 (Hex5Pen3, Fig. 1) in XyGOs from root tissues of wheat may suggest the presence of XyG of XXXG-type in these tissues. This ion was absent in XyGOs from wheat shoots. Although we were not able to identify unambiguously the presence of XXFG or XXLG structures characteristic of XyG of XXXG-type, our data is in agreement with a recent study showing that cell walls of young root tissues in rice contain XyG of XXXG-type, but not in shoot tissues [20]. Unfortunately, our ESI-MS instrument is not sensitive enough to detect low amounts of fucosylated XyGOs, however monosaccharide analysis and GC-MS strongly support the presence of Fuc, Gal, and Ara in XyGOs from wheat roots (data not shown).

To further confirm the presence of fucosylated XyG in wheat, we sought to determine whether wheat roots contain a XyG-fucosyltransferase activity responsible for the incorporation of Fuc into XyG in their tissues. Our data indicate that detergent-solubilized proteins from root tissues have the capacity to transfer radiolabeled [^14^C]Fuc from GDP-[^14^C]Fuc onto tamarind XyG *in vitro*. Shoot tissues did not show any detectable activity. This result was expected, as fucosylated XyG was detected in roots and not in shoots (under our conditions). It is possible that shoots may have “F” side chain, but their amounts must be below detection limit for our instrument. A recent work in rice identified a XyG-fucosyltransferase gene (*Os02g0764200, OsMUR2*) through functional complementation approach [20]. However, the rice XyG-fucosyltransferase activity has not been characterized biochemically. Wheat XyG-fucosyltransferase described here shares biochemical characteristics with activity from pea including no requirement for divalent cations, enhancement of activity with addition of MgCl_2_ and CaCl_2_ and no enhancement of activity with the addition of MnCl_2_ [29]. These similar biochemical attributes suggest that these activities are the same and that the wheat XyG-fucosyltransferase is encoded by a member of the same GT family (GT37 family in the CAZy database, which is a plant-specific GT family [58]). GT37 family also includes two Arabidopsis α(1,2)fucosyltransferases specific for arabinogalactan-proteins (AGPs), AtFUT4 and AtFUT6 [59]. In Arabidopsis, a single gene, *AtFUT1*, is responsible for XyG fucosylation, as the *mur2* mutant that has a lesion in the *AtFUT1* gene completely lacks fucosylated XyG [60]. Currently, there is no genetic evidence to demonstrate that *OsMUR2* is a single gene responsible for XyG fucosylation in rice. According to eFP platform (http://bar.utoronto.ca/efprice/cgi-bin/efpWeb.cgi), the expression of *Os06g0212100* is also confined to roots in rice, similar to *OsMUR2* (Fig. 6), which may suggest more than one XyG-FUT genes exist in rice. The ortholog of *Os06g0212100* in wheat (*TaFUT-G*) has very low expression levels in roots (Fig. 5B).

**Figure 6:**
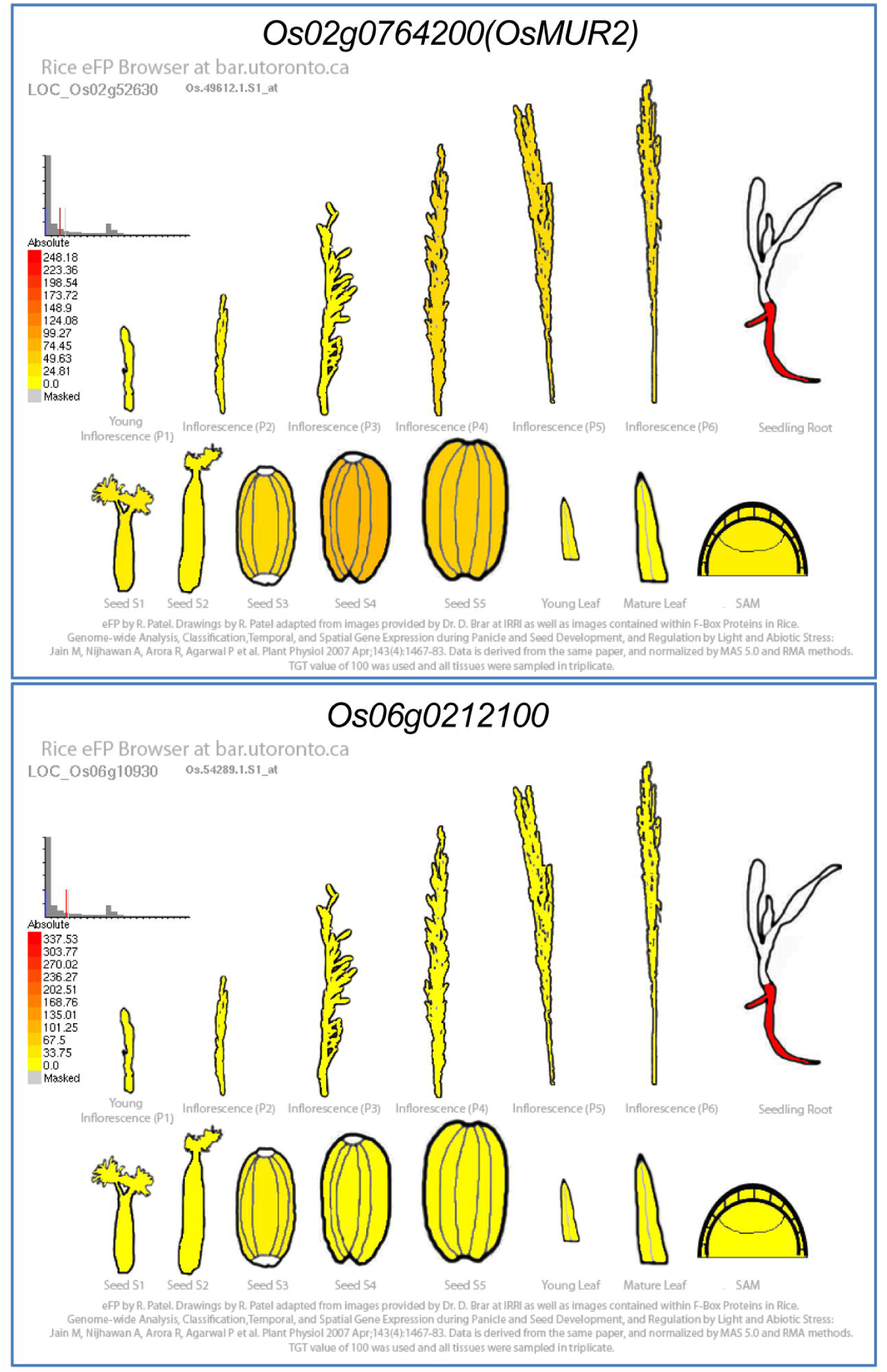
Expression profiles of *OsMUR2* and its closest homolog *Os06g0212100*. Expression data is obtained from eFP platform (http://bar.utoronto.ca/efprice/cgi-bin/efpWeb.cgi). Note that both genes have similar expression levels in roots and that the expression is mostly confined to roots.

Despite the identification of wheat XyG-fucosyltransferase activity, phylogenetic and expression profile analyses were not sufficient to identify true wheat orthologous proteins to AtFUT1 and PsFUT1. By the time this work was completed, the rice XyG-fucosyltransferase (Os02g0764200, OsMUR2) was not identified yet. At that time the wheat gene encoding for TaFUT-O was the most logical and promising candidate, as it is phylogenetically the most closely related to AtFUT1 and PsFUT1 (located in the same branch, Fig. 5A). However, *TaFUT-O* is expressed in both roots and shoots at similar levels, which does not fit with the fact that fucosylated XyG was mostly detected in roots and no detectable activity could be measured in shoots. According to expression data, the next most promising candidates are *TaFUT-F* and *TaFUT-I*, as they were expressed more highly in wheat roots than in shoots (Fig. 5B). Since we did not identify *TaFUT-I* through our strategy (sequence quality was low and was eliminated), we focused on *TaFUT-F*, as a second potential candidate. Unfortunately, none of these wheat FUTs showed *in vitro* XyG transfer activity. The recent early release of species at Phytozome 12 allowed the identification of *TaFUT-I*, which is the ortholog gene of *OsMUR2* in wheat. The failure of phylogenetic analysis in identifying the genuine wheat ortholog of AtFUT1 and PsFUT1 underscores the limitations of bioinformatics approach in identifying orthologs in protein families having a high degree of homology, such as the GT37 family. FUTs within a given plant tend to cluster more closely with each other than with FUTs from other plants. Thus, the only ways to identify functional orthologs is by heterologous expression/activity assays or by mutant complementation. It is challenging to select promising wheat candidates for functional analysis based only on amino acid sequence similarity to AtFUT1 and PsFUT1. For example, Os02g0764400 was previously identified as the only AtFUT1 ortholog in rice based on amino acid sequence similarity [61]. Overexpression of *Os02g0764400* in Arabidopsis *axy2.2* mutant did not restore XyG fucosylation. TaFUT-O is the closest homolog to Os02g0764400 in wheat (Fig. 5A), and our results also showed that TaFUT-O lacks XyG-fucosyltransferase activity, which is consistent with genetic complementation data from *Os02g0764400*.

The presence of Fuc residues in root XyG and less in coleoptile XyG is puzzling. One possible explanation could be the physiological role of Fuc residues in root interactions with microorganisms in soil. Several plant pathogens such as *Ralstonia solanacearum* and *Pseudomonas aeruginosa* produce L-Fuc-binding lectins. Interestingly, the lectin from *Ralstonia solanacearum* showed a strong affinity toward fucosylated XyG of XXXG-type [62]. Knockdown of the *PsFUT1* transcript in pea roots results in a phenotype of wrinkled and collapsed cells visible through SEM [63]. In grasses, a Fuc deficient mutant or XyG-fucosylatranferase gene knockout is not available for comparison, and it is unknown if grasses conserved the ability to substitute Fuc with L-Gal. The possible explanation of non-detection of fucosylation of XyG in wheat coleoptile using our ESI-MS instrument could be that it occurs at specific developmental stages and Fuc is removed by a fucosidase activity in later developmental stages. We showed that XyG-fucosyltransferase activity was higher in 5-day old seedlings and decreased drastically at 8 days of growth. No attempts were made to analyze XyG from roots of 8-day old wheat seedling to determine whether fucosylation of XyG occured at early developmental stages is maintained in walls of older seedling. Tissue-specific differences in XyG fucosylation are not unusual in plants. For example, certain species within the *Poaceae* appear to have phloem cells containing fucosylated XyG, while neighboring cells do not [18]. Also in the *Solanaceae*, fucosylated XyG of XXXG-type has been observed specifically in pollen tubes [64].

In conclusion, our present work demonstrates that etiolated wheat seedlings contain the machinery necessary to produce fucosylated and non-fucosylated XyGs (both XXXG-type and XXGGG-type). Although the identification of wheat XyG-fucosyltransferase activity may suggest that the XyG biosynthetic mechanism may be conserved in both monocots and dicots, biochemical and enzymological demonstration is still lacking and more experimental work is needed to elucidate XyG biosynthetic mechanism in monocots. It is currently not known if the “xylosyl-glucose” backbone synthesis in grasses occurs in the Golgi and involves concomitant Glc and Xyl incorporation by XyG-xylosyltransferase and XyG-glucan synthase activities, as described in dicots [26–29]. Such a cooperative mechanism has yet to be demonstrated in grasses.

## Materials and Methods

### Plant Material and chemicals

Winter wheat (*Triticum aestivum* L.) seeds were grown hydroponically in the dark for 6 d at 24°C using DynaGro 7-7-7 plant fertilizer (Richmond, CA). UDP-[^14^C]Xyl (9.78 GBq.mmol^−1^) and GDP-[^14^C]Fuc (7.4 GBq.mmol^−1^) were obtained from NEN (PerkinElmer, Boston, MA). The UDP-Glc, Dowex 1X80-100 (Cl^−^) ion-exchange resin, Sepharose-CL-6B and all the other chemicals were purchased from Sigma (St Louis, MO). Bio-gel P2 was from Bio-Rad. Purified pea cell wall XyG was a gift from Dr. Gordon Maclachlan (McGill University, Montreal, Canada). XyG-specific endo-β-(1,4)-glucanase (XG5, EGII, from *Aspergillus aculeatus*, 336 units/mg powder) was generously provided by Novozyme (Denmark). Tamarind XyG, rhamnogalacturonans-I (RGI, from soybean and potato), pectic galactan (from potato), arabinan (from sugar beet), AZO-xylan (Birchwood), AZO-carob galactomannan, AZO-wheat arabinoxylan, and AZO-carboxymethyl-cellulose were purchased from Megazyme International (Bray, Ireland). Oligonucleotide primers and TOP10 DH5α competent *E. coli* were from Invitrogen (Carlsbad, CA). Taq DNA polymerase, Q5 DNA polymerase, and restriction endonucleases were from New England Biolabs (Ipswich, MA). Pfu Ultra II fusion polymerase was from Agilent (Santa Clara, CA). Kapa HiFi HotStart polymerase was from Kapa Biosystems (Boston, MA). Spin columns for plasmid purification were from Epoch Biolabs (Sugar Land, TX) using the miniprep solutions and protocol from Qiagen (Valencia, CA). Gel purification was conducted using the Wizard SV gel and PCR cleanup system from Promega (Madison, WI). All primers used in this study are listed in Table 5.

**Table 5:**
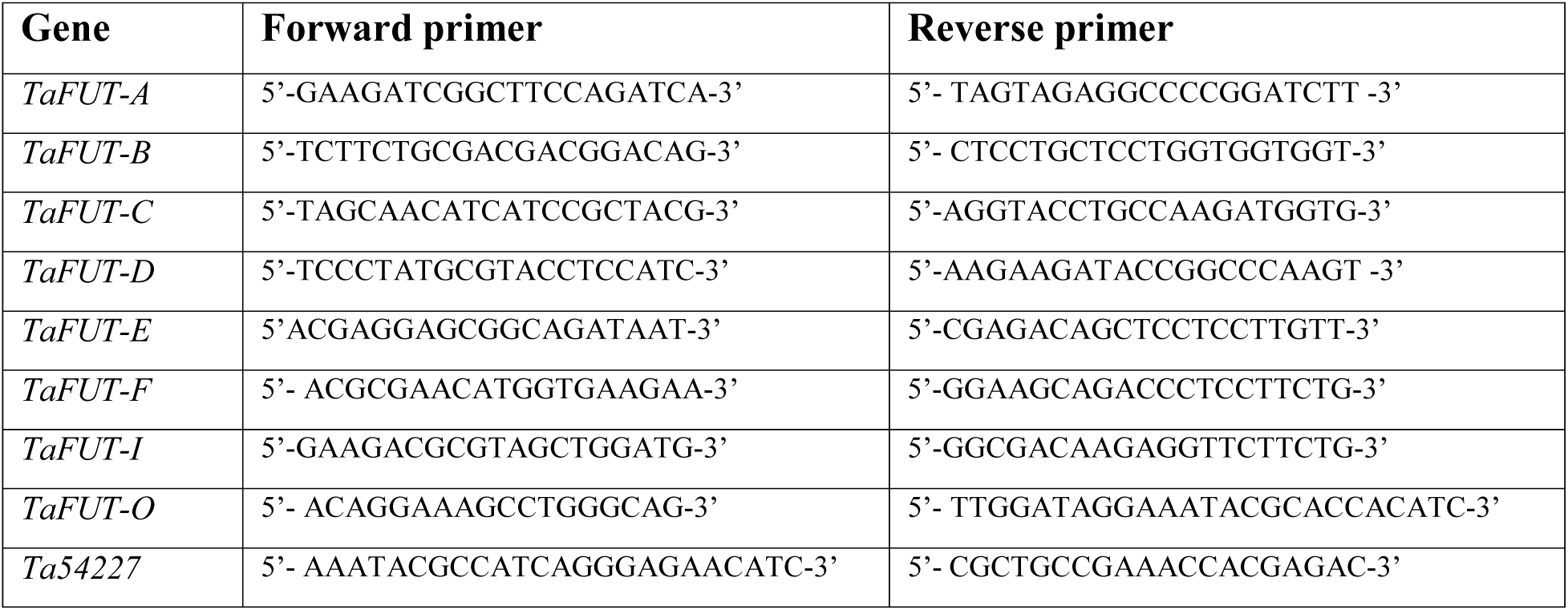
List of primers used for expression profiling of wheat *FUT* genes by RT-PCR.

### Purification of XyG-specific endo-β-1,4-glucanase XG5

The purification of XyG-specific endo-β-1,4-glucanase (XG5) was carried out according to [40] with some modifications. The crude XG5 provided by Novozyme was dialyzed against 50mM acetate buffer (pH 5.0) for two days in dialysis membrane having a MW cut off of 12-14kDa (Spectrum, Houston, TX). Dialyzed filtrate was applied to a Sep-Pak^®^ Vac Accell Plus QMA anion-exchange disposal column, 1cc/100mg, 37-55µm (Waters, Milford, MA) and the activity eluted with 50mM acetate buffer (pH 5). Fractions (0.5ml each) with high endo-β-1,4-glucanase activity were pooled and fractionated by gel permeation chromatography on a 1×30cm Superose 12 (12/300 GL) column (GE healthcare, Pittsburgh, PA). The column was eluted with 200mM sodium phosphate buffer (pH7) at a flow rate of 0.2ml/min and fractions collected every 2min. Active fractions were pooled, diluted to 1-5 units/µl, and stored at −20°C until use. Purified XG5 was tested for contaminating hydrolase activities on AZO-xylan (Birchwood), AZO-carob galactomannan, AZO-wheat arabinoxylan, and AZO-carboxymethyl-cellulose, and directly on tamarind XyG and screened for released monosaccharides. Also purified XG5 was tested for contaminant glycosidase activities on several paraphenyl-glycosides and. Under our conditions, no contaminant hydrolase and glycosidase activities were detected. However, when XG5 was tested on MLG according to [65], we detected mild β-glucan hydrolase activity, which releases mainly two diagnostic oligosaccharides, the tri-saccharide G-(1,4)-G-(1,3)-Gr and the tetra-saccharide G-(1,4)-G-(1,4)-G-(1,3)-Gr (in which G is β-D-Glc*p* and Gr is the reducing terminal residue). The release of these MLG oligosaccharides were estimated by the phenol-sulfuric acid method [66].

### Golgi-enriched microsomal membranes preparation

Tissues from 6-day old etiolated wheat and pea seedlings (grown on vermiculate) were used for microsomal preparation as described earlier [67] with minor modifications. Briefly, ∼25g of tissues were harvested and ground with a mortar and pestle in 50mL of extraction buffer (0.1M HEPES-KOH pH7, 0.4M sucrose, 1mM DTT, 5mM MgCl_2_, 5mM MnCl_2_, 1mM phenylmethylsulphonyl fluoride, 1 tablet of Roche complete protease inhibitor cocktail). The suspension was filtered through two layers of miracloth, and the filtrate centrifuged at 3,000x*g* for 20min. The residual, ground tissue was kept at −20°C for cell wall preparation and XyG extraction (see below). The resulting supernatant was layered over 1.8M sucrose cushion buffer and centrifuged at 100,000x*g* for 60min. Total microsomal membranes located at the top of the 1.8M sucrose cushion were collected and pelleted by centrifugation at 100,000x*g* for 30min. The pellet was resuspended in 200µl extraction buffer and stored at −80°C until use. This standard procedure usually yields membrane fractions with protein concentration of ∼5-6µg/μL. Protein content was estimated using the Bradford Reagent (Sigma) and various concentrations of bovine serum albumin (BSA) as standards.

### XyG-fucosyltransferase assays

#### Standard XyG-fucosyltransferase assay

The activity was measured as described earlier [44] with minor modifications. The reaction mixture (70µL final volume) contained the Triton X-100-solubilized enzyme from microsomes of 6-day-old wheat seedlings (40µl containing ∼0.2mg protein), 100µg tamarind XyG, and 6µM GDP-[^14^C]Fuc (65,000 cpm) or 30nM GDP-[^3^H]Fuc (70,000 cpm). Reactions were incubated for 2h at room temperature and terminated by adding 1mL of cold 70% (v/v) ethanol and precipitated for at least 1h at −20°C. Insoluble products were pelleted by centrifugation at 10,000x*g* (10min, 4°C), washed three times with 1mL of cold 70% (v/v) ethanol to remove excess GDP-[^14^C]Fuc, and the incorporation of [^14^C]Fuc into pellets (cpm/reaction) was determined by scintillation counting using a Beckman Coulter LS 6500 counter.

#### Effect of pH and divalent ions on wheat XyG-fucosyltransferase

The effect of pH on wheat activity was tested using Triton X-100 extracts under the same conditions as above but using 0.1M MES-KOH buffer for pH 4 and 5, and 0.1M HEPES-KOH buffer for pH 6 and 8. The effect of 5mM divalent ions (MnCl_2_, MgCl_2_, CuCl_2_, NiCl_2_, and ZnSO_4_) on wheat activity was tested under the same conditions as above, except that Golgi-enriched microsomes used were prepared with extraction buffer lacking any ions. Radiolabeled Fuc incorporation into pellets (cpm/reaction) was determined as described above.

### Preparation of XyG oligosaccharides (XyGOs)

Lyophilized KOH extracts (8mg) were resuspended in 0.4mL of 20mM sodium acetate buffer, pH5, to which 40 units of purified XyG-specific endo-β-(1,4)-glucanase (XG5) were added. The reaction mixtures were incubated at 37°C for 16 h with stirring. To terminate the reactions, mixtures were boiled (10min) and polymers were precipitated with 70% (v/v) cold ethanol for 1 h at −20°C. After centrifugation (14,000x*g*, 10min), the supernatant was collected and ethanol removed by evaporation at 60°C. Control reactions containing XG5 enzyme were boiled immediately after adding the enzyme and then incubated at 37°C for 16 h. Released XyGOs were used directly in ESI-MS and CID-MS/MS analyses.

### Electrospray ionization mass spectrometry (ESI-MS) and collision induced dissociation (CID)-MS/MS analyses

Samples (wheat XyGOs) containing 0.5mg carbohydrate were dissolved in 50% (v/v) methanol containing 0.1% (v/v) acetic acid. The sample was injected into the ESI source at a rate of 3µL/min using a Cole-Parmer 74900 syringe pump. ESI-MS spectra were acquired on an Esquire 6000 Ion Trap analyzer (Bruker Daltonics, Bremen, Germany) operated in positive ion mode with a capillary voltage 4kV, drying gas temperature 300°C, drying gas flow rate 5L/min and nebulizer pressure 10 psi. Nitrogen was used as both the nebulizing gas and drying gas. The mass range scanned was from 200 to 1500 atomic mass units. For CID-MS/MS, the spectra were recorded using the same instrument; the parent ions ([M+Na] ^+^) were selected in the first MS spectrum, then MS/MS spectra of daughter ions were obtained using collision induced dissociation (CID) with helium as the collision gas (introduced into the system to an estimated pressure of 4 x10^−6^ mbar). The amplitude of the excitation was 1 V. Instrument control and data acquisition were performed with Esquire 5.0 software.

### High pH anion exchange chromatography (HPAEC) analysis

#### Monosaccharide composition

Xyloglucan oligosaccharides (1mg) were mixed with trifluoroacetic acid (TFA) to a final concentration of 2M in a glass vial sealed with Teflon lined cap and autoclaved for 1 h at 120°C. The hydrolysates were desalted with Bio-Rex MSZ 501 resin (Bio-Rad) before analysis by HPAEC on a CarboPac PA20 column (Thermo-Dionex). The column was eluted at a flow rate of 0.5mL/min by isocratic elution for 30 min with 2.5mM NaOH solution. Monosaccharides Fuc, Ara, Gal, Glc, and Xyl were used as standards. The standards were run before analysis of samples to make sure that their elution profiles did not change between injections.

#### Analysis of [^14^C]Fuc-labeled tamarind XyG oligosaccharides

[^14^C]radiolabeled tamarind XyGOs were prepared by digestion of [^14^C]Fuc-tamarind XyG (produced by wheat fucosyltransferase reactions) with endoglucanase from *Trichoderma* sp. (E-CELTR, Megazyme, Bray, Ireland). [^14^C]Fuc-tamarind XyGOs were desalted with Dowex before analysis by HPAEC on a CarboPac PA200 column (Thermo-Dionex) as described earlier [29] with some modifications. Briefly, the column was eluted at a flow rate of 0.5mL/min for 60min at 30°C with an isocratic solution of 100mM NaOH containing 22mM sodium acetate; [^14^C]radiolabeled samples were injected in several repetitions to collect enough material for cpm counting. Known XyGO standards (*i.e.*, XXXG, XXFG, XLFG, XLLG, XLXG/XXLG) were included with [^14^C]radiolabeled samples as a control to make sure that their elution profiles did not change between injections.

### Identification of putative wheat fucosyltransferase (FUT) genes and phylogenetic analysis

Putative wheat FUTs were identified from EST databases, sequence reads from the 5X wheat genome (http://www.cerealsdb.uk.net/), and the partially assembled 5X wheat genome [68]. We developed two *in-house* scripts: one was used to screen wheat transcriptomes and the second for phylogenetic analysis. Both scripts were described in [69]. Briefly, these scripts perform a tBlastn (NCBI) search using rice and Arabidopsis GT protein sequences as queries and collect all the hits (E<0.05). After removal of redundancy (same accession number), the ESTs were merged into unique contigs using the CAP3 assembly program [70]. For the identification of contigs corresponding to various regions of the same gene sequence, it was necessary to manually perform several alignments using the ClustalW program and compare similarities with the closest full-length rice gene sequences. This step reduced the number of candidate wheat sequences and allowed the identification of start and stop codons of the wheat FUT genes. Additional wheat GT37 members were identified from the Roche 454 wheat genome sequence (http://www.cerealsdb.co.uk) using wheat FUT nucleotide sequences as a query. Obtained sequences were assembled using the built-in CAP3 program. Assembly and analysis of the 5X wheat genome allowed for the identification of additional putative wheat FUTs using Brachypodium gene identifiers as a query ([68]; http://mips.helmholtz-muenchen.de/plant/wheat/uk454survey/index.jsp). More wheat FUT sequences were found at Phytozome v12.1.6 (the Plant Comparative Genomics portal of the Department of Energy’s Joint Genome Institute), under early Release Species, (https://phytozome.jgi.doe.gov/pz/portal.html#!search?show=BLAST&method=Org_Taestivum_er). All hits above or equal to the score 920 (E≤2.5e-83) were collected. The final list of sequences was checked for the presence of the FUT domain using the CCD program at NCBI through the BLASTp program [71]. Rice GT37 members in the CAZy database were downloaded from GenBank. Putative full-length FUT protein sequences from rice, Arabidopsis, and wheat were used in the phylogenetic analysis along with partial sequences (ESTs or contigs) from wheat. Protein sequences were aligned using Muscle multiple sequence alignment [72]. Phylogenetic analysis was performed in the Phylogeny.fr platform (http://www.phylogeny.fr/[73]) using ‘One Click’ mode, which uses pipeline chaining of the following programs MUSCLE for multiple alignment [72], PhyML for tree building [74], and TreeDyn for tree rendering [75]. Default parameters were used for the phylogenetic analysis including Maximum-likelihood tree construction to infer phylogenies, which is commonly accepted as the most accurate approach in molecular phylogenetics [76]. PhyML is run with the aLRT statistical test [77] of branch support (to infer bootstrap values) is based on an approximation of the standard Likelihood Ratio Test. The number of bootstrap replicates is limited to 100. However, to confirm the stability of the phylogenetic results, we run ‘A la Carte’ mode using several different programs with various options. All methods output the same trees with branch supports that are highly correlated.

### Cloning of TaFUT-F and TaFUT-O genes

The original EST designated TaFUT-F (BU099714, full tillering stage drought stressed cDNA library from the USDA) was 667bp long and missing sequences from both the 3’ and 5’ ends. The 3’ end sequence and 3’ untranslated region (UTR) of *TaFUT-F* were verified by 3’ Rapid Amplification of cDNA ends (RACE) using the GeneRacer kit (Invitrogen). Two micrograms of total RNA from wheat roots was reverse transcribed using SuperScriptIII reverse transcriptase with the oligo(dT) primer including an adaptor sequence included with the kit. First round PCR was performed with a forward primer in the EST (5’-GCGGAGATATATCTGCTCAGCCTC-3’) and the oligo(dT) adaptor primer using 2µL of cDNA template. Touchdown (TD)-PCR was performed using with the following program: 96°C for 3min; 20 cycles of 96°C for 15s, 65°C for 15s decreasing by 0.5°C each cycle, 72°C for 90s; 20 cycles of 96°C for 15s, 55°C for 15s, 72 for 90s; and final extension at 72°C for 5 min. Second round PCR was performed using 1µL of the 1st round PCR product as template, a forward nested primer in the EST (5’-GTGATGTTCAAGCCGGACA-3’) and the nested 3’ primer included in the kit. Cycling conditions were the same as first round PCR. The 5’ end was cloned through RNA-ligase mediated RACE (RLM-RACE) using the GeneRacer kit (Invitrogen). Total RNA template was made according to manufacturer’s instructions. Reverse transcription was conducted as for 3’ RACE. First round PCR was conducted on the reverse transcribed RNA using the GeneRacer 5’ forward primer and a reverse target specific primer (TSP) located in the EST (5’-AATGGCTCGCAGAACAGCTC-3’). Touchdown-PCR was conducted using Kapa HiFi polymerase and the same program as above except the annealing temperature started at 72°C and touched down to 62°C. Nested PCR was performed using 1µL of the 1^st^ round PCR product as template and the 5’ GeneRacer nested primer and a nested TSP (5’-ACATGGCGGACTGGTACCTGCTGTG-3’) using the same PCR program. The full-length *TaFUT-F* cDNA was then cloned from cDNA generated for 5’ RACE using forward primer (5’-ATGCTGCGACGGGACGTC-3’) and reverse primer (5’-GCACACGCCACCAGGTTTC-3’) in a 50µL reaction containing 2µL cDNA template, 5µL 10X reaction buffer, 250µM each dNTP, 0.5µM each primer, and 1µL Pfu Ultra II Fusion DNA polymerase using the following program: 94°C for 3min; 30 cycles of 94°C for 30s, 57°C for 30s, 72°C for 90s; and a final extension at 72°C for 5 min.

For *TaFUT-O*, the second exon was found by searching the wheat genome with the gene identifier “*Bradi3g58030.*1” ([68], http://mips.helmholtzmuenchen.de/plant/wheat/uk454survey/index.jsp). Primers were designed specifically from the entry “Traes_Bradi3g58030.1_000002_B.” The full-length *TaFUT-O* transcript was amplified from cDNA prepared for 5’ RACE as a template (described above) using a forward primer designed from Brachypodium homolog, *Bradi3g58030*, 5’-ATGGACCTCAAGGAGCGGATCC-3’, and a reverse primer (designed in the 3’UTR) 5’-TTGGATAGGAAATACGCACCACATC-3’ in a 25µL PCR reaction containing 2µL cDNA template, 0.8µM each primer, 250µM each dNTP, 5µL of 5X Q5 buffer, and 1U of Q5 Hot-start DNA polymerase. Touchdown PCR was conducted with the following program: 98°C for 30s; 20 cycles of 98°C for 10s, 72°C for 20s decreasing by 0.5°C each cycle, 72°C for 1 min; 20 cycles of 98°C for 10s, 62°C for 20s, 72 for 1 min; and final extension at 72°C for 5 min.

*TaFUT-D* was cloned from cDNA produced from wheat root RNA (prepared as described above for 3’ RACE) using Kapa HiFi HotStart polymerase. The reaction was set up according to manufacturer’s instructions using forward primer (5’-ATGGGGAGGAGCGGCG-3’) and reverse primer (5’-CAGCTACAACCGAGTTTCATTT-3’) and the following PCR program: 95°C for 3 min; 30 cycles of 98°C for 20s, 65°C for 15s, 72°C for 60s; and a final extension at 72°C for 3 min.

### Expression profiling of wheat FUT genes

Total RNA was isolated from the roots and coleoptiles of etiolated wheat seedlings using Direct-zol RNA Miniprep (Zymo Research, Irvine, CA). Two micrograms of total RNA was reverse transcribed with an oligo(dT) primer using the Applied Biosystems High Capacity cDNA Reverse Transcription kit according to the manufacturer’s instructions. Genomic DNA was extracted from shoot tissues using a CTAB extraction method [78]. Complementary DNA (cDNA) from roots and shoots of etiolated wheat was used for PCR screening of nine *TaFUT* genes. Gene-specific primers were designed to amplify products from these *TaFUT* genes ranging in size from 220-398bp (Table 5). The products were confirmed through sequencing. Template cDNA was prepared as described in the previous section. Products were amplified in a 20µL PCR reaction containing 1-2µL cDNA template (normalized using control primers), 1µM each primer, 250mM each dNTP, 2µL standard Taq buffer, and 1U *Taq* DNA polymerase. Cycling conditions were as follows: 96°C for 2min; 25 cycles of 96°C for 15s, 58°C for 15s, 72°C for 40s; and a final extension at 72°C for 7min. The annealing temperatures for *TaFUT-O* and Ta54227 were 64°C and 70°C, respectively. Control primers were for *Ta54227*, a gene encoding a cell division control protein that is a member of the AAA-superfamily of ATPases [79]. It was chosen as a control based on its stable expression between different tissue types and outperformance of common controls such as actin or ubiquitin [79].

### Expression in Pichia pastoris cells

Transformation of *Pichia pastoris* was carried out according to the manufacturer’s manual (Invitrogen) with minor modifications. A colony of *Pichia pastoris* strain X-33 was picked from a Yeast Peptone Dextrose [YPD; 1% (w/v) yeast extract, 2% (w/v) peptone, 2% (w/v) agar] plate and grown in 5mL liquid YPD in a 50mL conical tube overnight at 28°C with constant shaking (220 rpm). Five-hundred microliters of this culture was used to inoculate 250mL fresh YPD in a 1L flask. Cells were grown to an OD600 ∼1 (∼10 hrs) and harvested by centrifugation at 3000x*g* for 5 min. The pellet was re-suspended in 200mL ice-cold sterile water and centrifuged. The pellet was then re-suspended in 100mL ice-cold sterile water and centrifuged again. The pellet was re-suspended in 10mL of ice cold 1M sorbitol and centrifuged again, and the final pellet was resuspended in 300µL 1M sorbitol. Competent cells were kept on ice and used for electroporation the same day.

Constructs of His-tagged FUT genes in pPICZa were prepared for electroporation as follows. A single construct was grown overnight in 18mL LB (6 × 3mL cultures) containing 100 μg/mL Zeocin, and bacteria cells were used for plasmid DNA preparation using the standard miniprep procedure (Qiagen). The purified plasmid was precipitated by adding 1/10^th^ volume 3M sodium acetate and 2.5 volume 100% cold ethanol, left overnight at −20°C, recovered by centrifugation at 14,000x*g* for 20 min at 4°C, washed once with 200µL cold 70% ethanol, and re-suspended in 15µL (typical yield is ∼17μg of DNA). Ten micrograms of the plasmid was linearized with 10U of *Pme*I in a 10µL volume, and 1µL was used to monitor the status of the linearization on a DNA gel.

Pichia competent cells (80µL) were combined with the linearized plasmid and allowed to incubate on ice for five minutes prior to electroporation. The cells were pulsed at 1.8kV, 200 ohms, 25 µF in a Gene Pulser II Electroporation System (BioRad). Following electroporation, 1mL of 1M sorbitol was added to the electroporation cuvette and the contents were transferred to a 15mL conical tube and incubated for 2 h at 28°C. Cells were plated on YPDS [1% (w/v) yeast extract, 2% (w/v) peptone, 1M sorbitol, 2% (w/v) agar] containing 100-200µg/mL Zeocin. After 2-3 days of incubation, ten colonies were picked and grown in 3mL YPD containing 100µg/mL.

Two to five colonies for each construct were chosen for the production of the proteins. First, 100 µL of a colony’s glycerol stock was added to 15mL BMGY [1% (w/v) yeast extract, 2% (w/v) peptone, 100mM potassium phosphate buffer pH 6, 1.34% (w/v) yeast nitrogen base, 4×10^−5^% (w/v) biotin, 1% (v/v) glycerol] and grown at 28°C overnight with constant shaking (220 rpm) until the OD600 was between 2 and 6. To initiate the induction of protein production, Pichia cells were harvested by centrifugation at 3,000x*g* for 10 min and re-suspended to an OD600 of 1 in 30-50mL BMMY [1% (w/v) yeast extract, 2% (w/v) peptone, 100mM potassium phosphate buffer pH 6, 1.34% (w/v) yeast nitrogen base, 4×10^−5^% (w/v) biotin, 0.5% (v/v) methanol]. Methanol was added every 24 h to a final concentration of 0.5% (v/v) to maintain the induction. One mL was taken from the culture each day to monitor the level of protein expression using western blot analysis. After 4 days of induction, microsomes were prepared as described earlier [31] with minor modifications. Cells were pelleted by centrifugation at 3,000x*g* for 15 min, re-suspended in 10mL extraction buffer (EB) [100mM HEPES-KOH pH 7, 0.2M sucrose, 1mM dithiothreitol (DTT), 5mM MgCl_2_, 5mM MnCl_2_, 1mM phenylmethanesulphonylfluoride (PMSF)], pelleted again, and resuspended in 5mL EB containing 1x complete protease inhibitor (Roche) and 50 µL RPI protease inhibitor cocktail VI (concentrations in vial: 200mM AEBSF, HCl; 10mM Bestatin; 3mM E-64; 2mM Pepstatin A; 2mM Leupeptin; 500mM 1,10 phenanthroline). Transgenic Pichia cells were broken in 15mL conical tubes by adding ∼2mL glass beads (425-600 µm diameter) and vortexing seven times, 30 s each time. Tubes were inverted each time while vortexing and kept on ice at least 1.5 min between each vortexing. Broken cells were centrifuged at 3,000x*g* at 4°C for 15 min to collect beads and remove cell debris. The supernatant was centrifuged at 100,000x*g* for 1 h at 4°C to collect microsomal membranes, and the pellet was re-suspended in ∼400µL EB. This procedure yielded membrane fractions ranging from 1 to 5 µg/mL of protein as determined by a Bradford assay.

### GenBank accession numbers of cloned wheat FUT genes

The nucleotide sequences of *TaFUT-D, TaFUT-F, and TaFUT-O* have been assigned the following GenBank accession numbers: MN529249, MN529250, MN529251, respectively.

## Acknowledgments

The authors would like to thank Dr. Kirk Schnorr from Novozyme (Denmark) for generously providing XyG-specific endoglucanase XG5, EGII from *Aspergillus aculeatus*. We also thank Dr. Michael Hahn, Dr. Siva Pattathil, and Dr. Samuel Hazen for their precious comments on the manuscript. This work is dedicated to the memory of Prof. Gordon Maclachlan (McGill University, Montreal, Canada).

## Abbreviations

ESI: Electrospray ionization
CID: Collision-induced dissociation
GC-MS: Gas chromatography mass spectrometer
Glc: Glucose
Xyl: Xylose
Gal: Galactose
Ara: Arabinose
Fuc: Fucose
XyG: Xyloglucan
XyGOs: Xyloglucan oligosaccharides
TXyG: Tamarind xyloglucan
HPAEC: High pH anion exchange chromatography
PAD: Pulsed-amperometric detection
AIR: Alcohol-insoluble residue
RT: Reverse transcription
CAZy: Carbohydrate-active enzyme

